# Diversity of reptile sex chromosome evolution revealed by cytogenetic and linked-read sequencing

**DOI:** 10.1101/2021.10.13.462063

**Authors:** Zexian Zhu, Kazumi Matsubara, Foyez Shams, Jason Dobry, Erik Wapstra, Tony Gamble, Stephen D. Sarre, Arthur Georges, Jennifer A. Marshall Graves, Qi Zhou, Tariq Ezaz

**Author notes:** These authors contributed equally to the work (Zexian Zhu, Kazumi Matsubara). Current affiliation: Department of Environmental Biology, College of Bioscience and Biotechnology, Chubu University, 1200 Matsumoto-cho, Kasugai, Aichi 487-8501, Japan (Kazumi Matsubara). Correspondence should be address to Q. Z. and T. E.

## Abstract

Reptile sex determination is attracting much attention because the great diversity of sex-determination and dosage compensation mechanisms permits us to approach fundamental questions about sex chromosome turnover and evolution. However, reptile sex chromosome variation remains largely uncharacterized and no reptile master sex determination genes have yet been identified. Here we describe a powerful and cost-effective “chromosomics” approach, combining probes generated from the microdissected sex chromosomes with transcriptome sequencing to explore this diversity in non-model Australian reptiles with heteromorphic or cryptic sex chromosomes. We tested the pipeline on a turtle, a gecko, and a worm-lizard, and we also identified sequences located on sex chromosomes in a monitor lizard using linked-read sequencing. Genes identified on sex chromosomes were compared to the chicken genome to identify homologous regions among the four species. We identified candidate sex determining genes within these regions, including conserved vertebrate sex-determining genes *pdgfa, pdgfra amh* and *wt1*, and demonstrated their testis or ovary-specific expression. All four species showed gene-by-gene rather than chromosome-wide dosage compensation. Our results imply that reptile sex chromosomes originated by independent acquisition of sex-determining genes on different autosomes, as well as translocations between different ancestral macro- and micro-chromosomes. We discuss the evolutionary drivers of the slow differentiation, but rapid turnover, of reptile sex chromosomes.

## Introduction

Sex can be determined either by genes on specialized chromosomes (genetic sex determination, GSD) or by environmental factors (environmental sex determination, ESD). Much of our knowledge on sex chromosome evolution has come from studies of model organisms such as *Drosophila*, chicken and mammals (principally humans and mice), in which species master sex determining genes have been identified ^1^. Their heteromorphic sex chromosomes can be easily identified by cytogenetic observations because the male-specific Y chromosome, or the female-specific W chromosome is morphologically different from the X or Z chromosome. Sex chromosome differentiation occurs as the result of suppression of recombination, and is manifested by massive accumulation of massive transposable elements and inactivation or loss of genes ^2^. The sex chromosomes of many model vertebrate species have been evolutionarily stable for more than 100 million years, judging from the homology of the pair of sex chromosomes within their clade ^3, 4^.

However, in many reptiles, amphibians and fish, there are frequent transitions between different sex determination mechanisms ^5, 6, 7^. Reptiles represent an extraordinary variety of sex determining mechanisms, including GSD and TSD, XY and ZW systems with varying degrees of sex chromosome differentiation ^8^. However, we know little about reptile sex chromosomes and sex determining genes.

The evolutionary variety of vertebrate sex determining systems has long been recognized. Cytological observations and limited gene mapping data reveal that multiple transitions between ESD and GSD, and between XY and ZW sex chromosome systems, have occurred in reptiles ^5^, teleost fish ^6^ and anurans ^7^. However, despite this variety, extensive cytogenetic mapping of the reptile orthologues of genes that are located on sex chromosomes of model organisms (e.g., human and chicken) revealed a surprisingly frequent over-representation of particular ancestral autosomes or genomic regions ^9, 10, 11^.

With the development of long-read sequencing and Hi-C technologies, many genomic consortia (e.g., Vertebrate Genome Project ^12^ and the Earth Biogenome Project ^13^ aim to finish the complete genomes of most vertebrate species on earth in the next few years. However sequencing projects usually represent sex chromosomes poorly; either the homogametic sex (XX female or ZZ male) is sequenced, and the male-specific Y or female-specific W is ignored; or the heterogametic sex only is sequenced with poor representation of the X or Z, and there is great difficulty in assembling the repeat-rich Y or W ^14, 15^.

Here, we develop a cost-effective method to identify genes borne on sex chromosomes, combining microdissection of sex chromosomes and high-throughput sequencing, followed by PCR validation and assessment as candidate sex determining genes. A similar method was pioneered to identify novel genes on the Y chromosome of marsupials ^16^. We applied the method to four reptile species, revealing the great diversity of sex chromosomes, and their independent evolutionary origins.

We chose four Australian reptile species, a turtle and three lizard species to represent the variety of reptile sex determining systems (**Figure 1**).The Murray River turtle *Emydura macquarii* ^17^ (referred as ‘river turtle’ hereafter) has a cryptic XX/XY sex chromosome system in which minimally differentiated X and Y are macrochromosomes, whereas the pink-tailed worm lizard *Aprasia parapulchella* ^18^ (referred as ‘worm lizard’ hereafter) has a highly differentiated XX/XY sex chromosome system in which the X and Y are microchromosomes. The marbled gecko *Christinus marmoratus* ^19^ (referred as ‘marbled gecko’ hereafter) has a pair of ZZ/ZW sex chromosomes, in which the Z and W heteromorphy involves pericentric inversion, whereas the spiny-tailed monitor lizard *Varanus acanthurus* ^20^ (referred as ‘monitor lizard’ hereafter) has ZZ/ZW heteromorphic sex chromosomes in which Z and W chromosomes are minimally differentiated microchromosomes (**Figure 1**).

**Figure 1.**
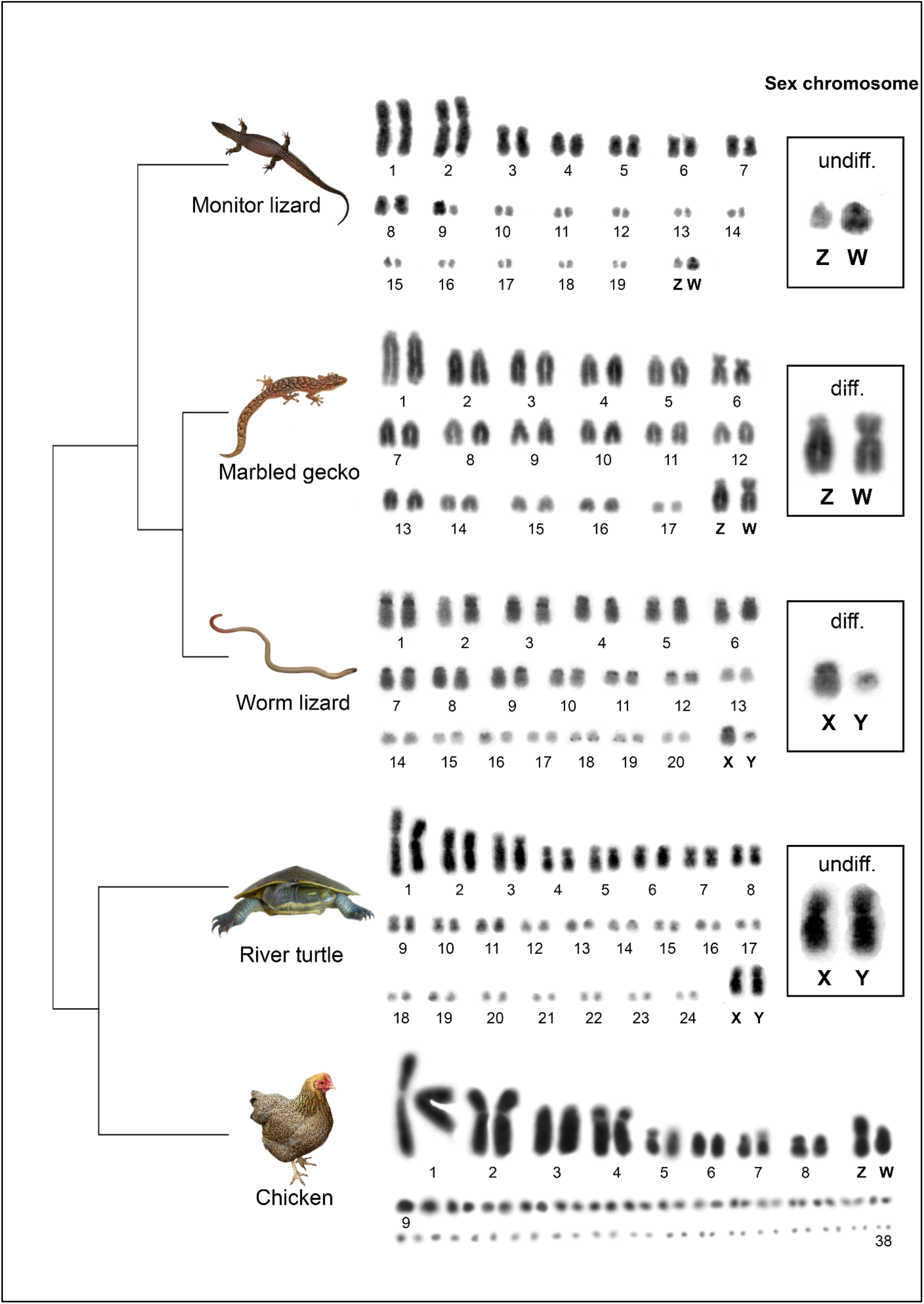
The diversity of reptile sex chromosomes. Cladogram and karyotypes of the studied reptile species river turtle (*Emydura macquarii*), worm lizard (*Aprasia parapulchella*), Marbled gecko (*Christinus marmoratus* and monitor lizard (*Varanus acanthurus*) with two types of sex chromosome systems. Species with differentiated sex chromosomes are labelled with “diff.”, otherwise with “undiff.”. Photo credit: see acknowledgements.

## Results

### Transcriptome and genome assemblies of sex chromosomes of the four reptile species

The four reptile species have cytologically distinguishable sex chromosome pairs (**Figure 1**, **Supplementary Fig. S1**); these were morphologically differentiated in the worm lizard and the monitor lizard, but subtle for the river turtle and the marbled gecko ^18, 19, 20^. For each species, we microdissected each of their sex chromosomes, performed linear genome amplification and validated the sex chromosome specificity of the DNA products by chromosome painting (**Figure 2A, Supplementary Fig. S1**).

**Figure 2.**
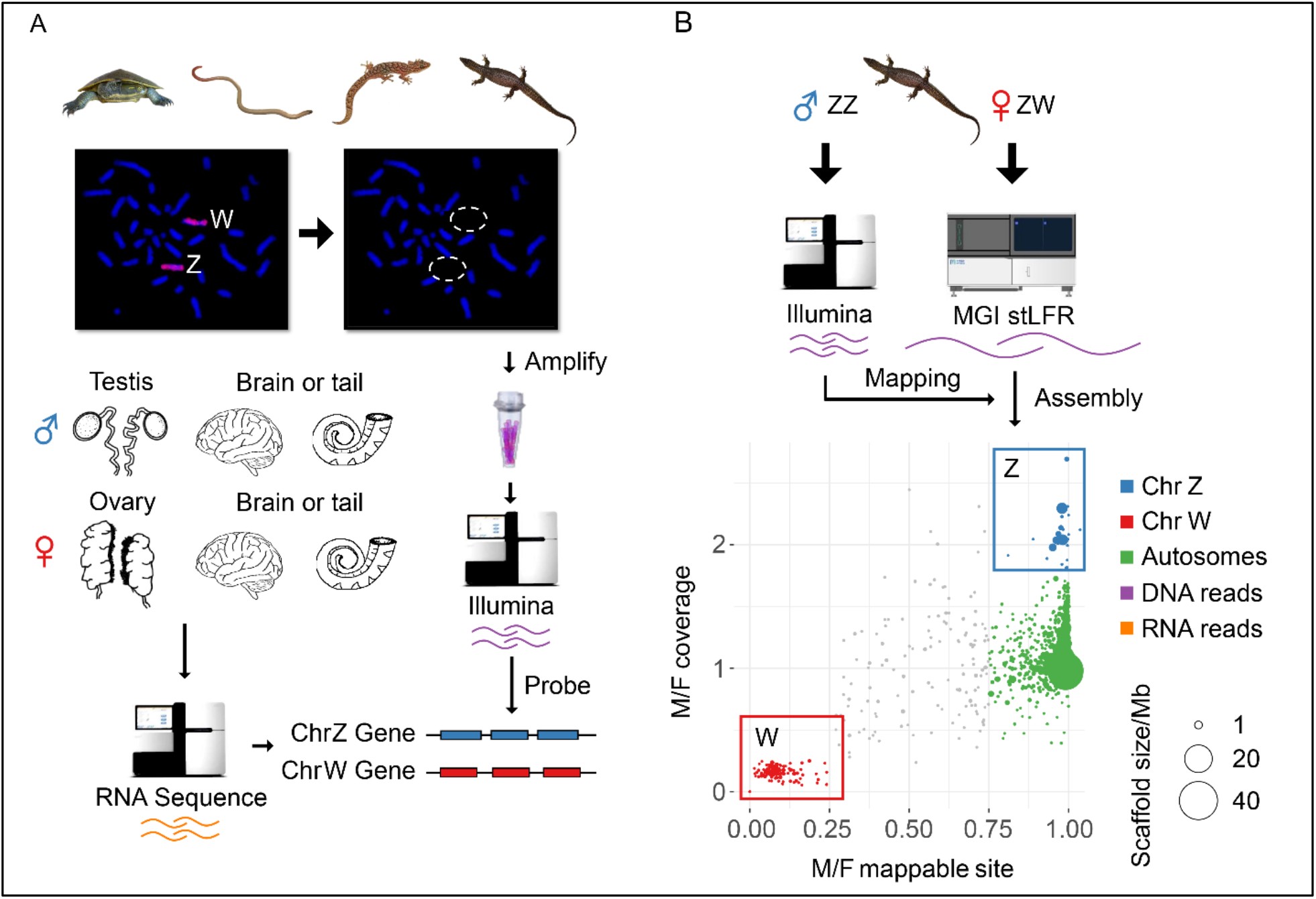
Transcriptome and genome assemblies of four Australian reptile species. **A**. The pipeline of identifying sex-linked transcripts of river turtle, worm lizard and marbled gecko using transcriptomes of sexed tissues, and amplified probes from the microdissected sex chromosomes. The probes have been validated by chromosome painting, and Illumina reads generated from the probes were used to identify the sex chromosome genes from the *de novo* assembled transcripts. **B**. Identifying sex-linked sequences of monitor lizard based on the *de novo* genome assembly generated from linked (stLFR) reads. For the assembled Z-linked sequences (in blue), we found a 2-fold male vs. female (M/F) ratios of Illumina DNA sequencing coverage, but an equal mappable site between sexes. While the W-linked sequences (in red) exhibited a female-specific pattern of read coverage ratio and mappable sites. The size of each dot is scaled to its scaffold length.

For each sex chromosome, we generated up to 2Gb clean paired-end (PE) Illumina reads from the microdissected sex chromosome DNA **(Supplementary Table S1)**. To identify genes borne on sex chromosomes, we also produced 2Gb transcriptomes from the gonads and brain tissues for males and females of monitor lizard, river turtle and marbled gecko and somatic transcriptomes (tail tissue) from a male and female worm lizard (**Figure 2B**). Genomic reads derived from each sex chromosome were then used to identify sex-linked genes from *de novo* assembled transcript sequences of each species. We annotated a total of 11299, 15202 and 10507 non-redundant transcripts, respectively for the worm lizard, the marbled gecko, and the river turtle, using chicken genes as a reference for each.

For the monitor lizard inferences based on transcripts and microdissected sex chromosome sequences were uncertain because its sex chromosomes were reported to have originated from translocations between fragments of multiple ancestral autosomes (also see below) ^21^, as well as the poor quality of sequences obtained from microdissected monitor lizard sex chromosomes. Therefore, we generated 200 Gb (135x genomic coverage) single-tube long fragment linked reads (stLFR)^22^ from a female individual, and 30 Gb Illumina PE reads from a male individual. We performed *de novo* genome assembly and produced a female draft genome with the total length of 1.46Gb and the scaffold N50 length of 12.8Mb (**Supplementary Table S2**). The high continuity of the draft genome was evident from 94 very large scaffolds that accounted for 80% of the entire genome. Using protein sequences of human and chicken as reference, we annotated a total of 14521 genes for the monitor lizard and identified its sex-linked sequences based on the comparisons of mapped read patterns between sexes. The putative W-linked scaffolds showed female specificity in both their mapped read number and mapped sites, whereas the Z-linked scaffolds showed a 2-fold increase of male mapped reads compared to that of female mapped reads, but an equal number of mapped sites for putative autosomal scaffolds between sexes (**Figure 2B**). Using this approach, we identified 10.81 Mb Z-linked scaffolds with 337 genes, and 7.10 Mb W-linked scaffolds with 87 genes.

### Identification of sex-linked genes

To identify the sex chromosome-borne genes in three species other than the monitor lizard, we developed a pipeline to separately assemble transcripts of genes that are X- or Y-borne (or Z- and W-borne) using the sexed transcriptomes (**Figure 3A**). In brief, we considered that transcripts that were assembled using pooled RNA-seq reads of both sexes and could not be aligned using the sex chromosome DNA probes were autosomal genes. Conversely, male RNA-seq reads in XY species that could not be aligned to the female transcripts were assembled into candidate Y-borne transcript sequences. Then by comparing the mapped read numbers of Y- or X-borne probes for each candidate Y-borne transcript or each transcript assembled from female RNA-seq, we were able to categorize them into the genes that were specific to the X or to Y chromosome, or were shared between X and Y. We also conducted the same process for the ZW marbled gecko but in reverse.

**Figure 3.**
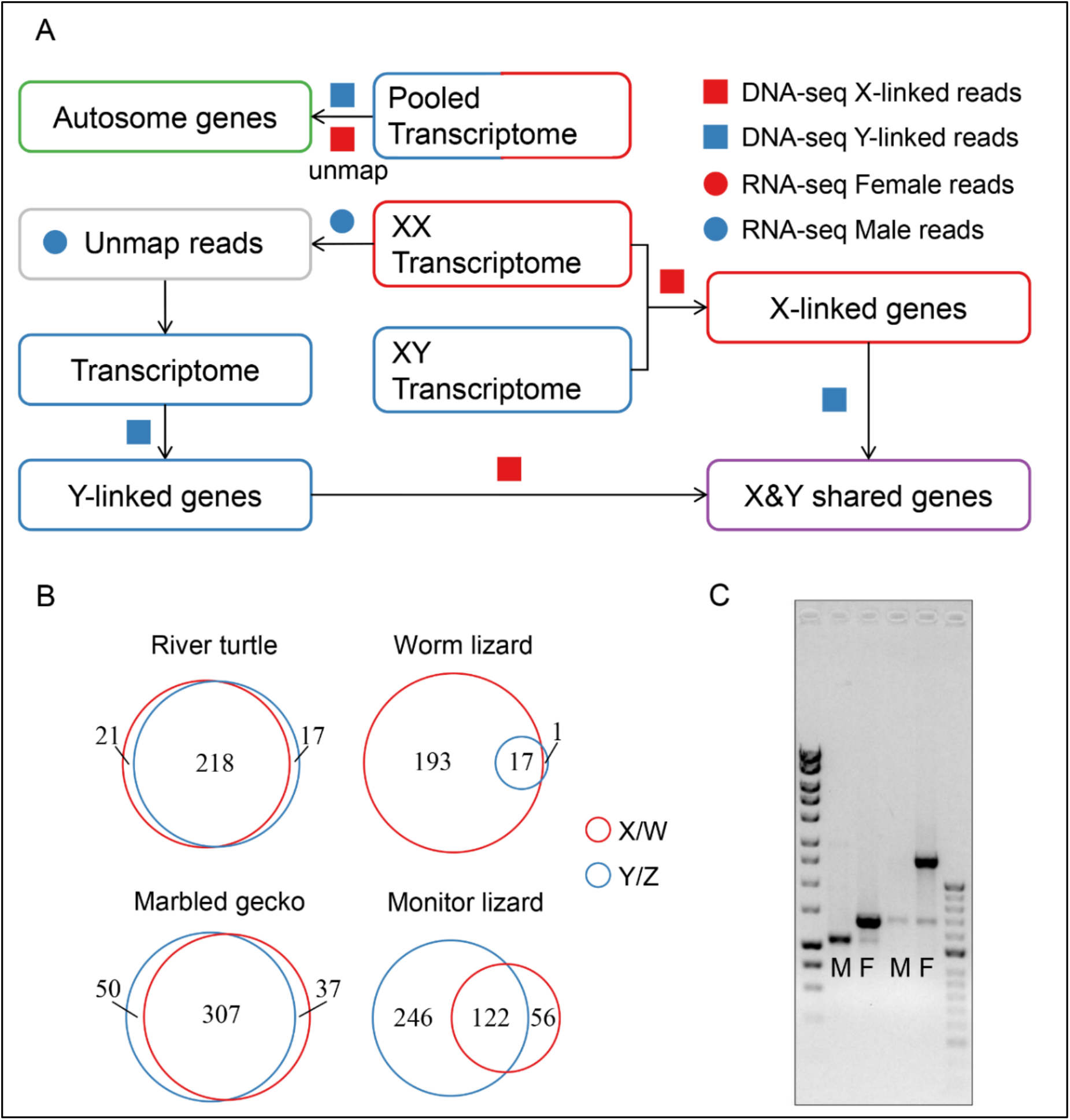
Processes involved in the identification of sex-linked genes. **A**. Pipeline to separately assemble the X-linked (red box), Y-linked (blue box) genes using the sexed transcriptomes (red: female; blue: male). Squares (DNA) and circles (RNA) refer to the reads from different resources. Shared genes, which can be aligned by probes from both sex chromosomes are labelled in purple colour. The autosomal genes, which cannot be aligned by the sex chromosome DNA probes are labelled in green; **B**. Numbers of sex-linked genes in the four reptile species. X-linked and W-linked genes are labelled in red colour, while Y-linked and Z-linked genes are labelled in blue colour. The overlapping areas refer to the genes shared between the two sex chromosomes; **C**. An example of PCR validation of sex-linked sequences of the monitor lizard. M refers male individual and F refers female individual; outside lane size standard 1kb (left) and 50bp (right) ladder.

Following our stringent filtering criteria (see Methods), we identified 193 X-borne genes, 1 Y-borne genes and 17 shared genes between X/Y chromosomes in the worm lizard, 21 X-borne genes, 7 Y-borne genes and 218 shared genes in the river turtle and 50 Z-borne genes 37 W-borne genes and 307 shared genes in the gecko (**Figure 3B, Supplementary Table S3**). We considered these numbers to be conservative estimates of the sex chromosome-borne genes in these species because genes with low expression levels may not be well assembled in our transcriptome data and there could also be a sampling bias in the sex chromosome probes captured by microdissection. We further designed primers spanning regions of insertions or deletions between the sex chromosomes and confirmed their length variations between sexes by PCR for the sex chromosome-borne genes of the monitor lizard (**Figure 3C**). We found no indel sequences within the coding regions of sex chromosome borne genes of the other three species, hence did not design primers for validation.

The proportion of genes that are specific for one or other sex chromosome, and the proportion that are shared, provide a good indication of the degree of genetic differentiation of the sex chromosome pair, and correlate well with our cytogenetic observations (**Supplementary Fig. S1**). The high numbers of genes shared between the X and Y chromosomes of the river turtle suggest that its sex chromosome pair is not highly differentiated, which is consistent with the subtle difference in size and morphology of the X and Y (**Figure 1**). Among the three lizard species, the numbers of shared versus sex chromosome-specific genes also implied different degrees of sex chromosome differentiation. Consistent with the cytogenetic data (**Figure 1, Supplementary Fig. S1**) ^18^, the Z and W chromosomes of the marbled gecko also shared most genes, whereas the monitor lizard showed an intermediate level of shared Z- and W-borne genes. In contrast, the worm lizard had many X-specific but very few Y-specific genes, and only a few X-Y shared genes, implying that the Y chromosome is highly degraded.

### Origins of sex chromosomes of the four reptile species

By mapping the orthologues of sex-linked genes of the four reptiles to the chicken genome (GGA), we found evidence for both independent origin and convergent evolution of sex chromosomes (**Figure 4, Supplementary Fig. S2**).

**Figure 4.**
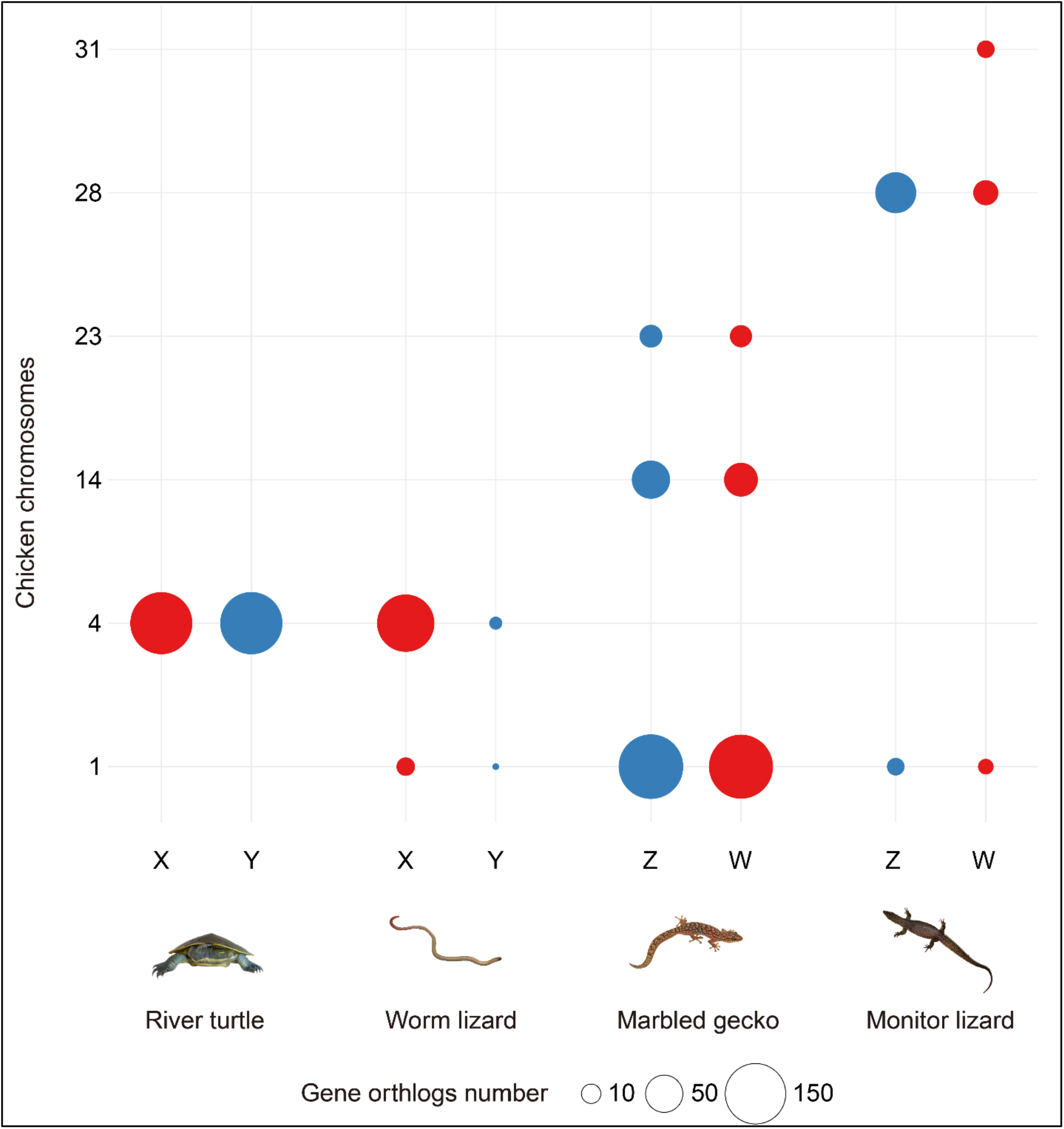
Independent origin of the sex chromosomes of four reptile species. The independent origins of the sex chromosomes of four reptile species with chicken chromosomes as reference. The bubble size is scaled to the number of orthologs of sex chromosome borne genes of each reptile species within the chicken genome, and the colour indicates the type of sex chromosomes (red for X and W, and blue for Y and Z).

In each species, genes borne on the sex chromosomes clustered together predominantly on a single chicken chromosome, though in three of the species there were other minor clusters. Sex chromosomes of the three lizards were homologous to quite different regions of the chicken genome, on chromosomes GGA1, GGA4 and GGA28 respectively, implying independent origins. However, the sex chromosomes of the river turtle largely overlapped with those of the worm lizard on GGA4q, the long arm of chicken chr4. This is unlikely to represent sex chromosome identity by descent, since the turtles are more closely related to birds (in which this region is autosomal) than they are to squamates, with divergence times of ~250 million years ago (MYA) and 285 MYA respectively ^23^.

Secondary sites of homology between our four reptile species and the chicken genome represent fragments of sex chromosomes with a different evolutionary origin. For instance, homologs of some river turtle sex chromosome-borne genes were found on chicken microchromosome GGA32 (**Supplementary Fig. 2**). This supports the hypothesis that the sex chromosomes of the river turtle originated by a recent translocation between an ancestral sex chromosome pair (GGA4) and a microchromosome pair ^17, 24^.

The sex chromosomes of marbled gecko were mainly homologous to GGA1p, with strong secondary signals on GGA14 and GGA23 that contained very similar numbers of Z- and W-borne genes. Genes on the sex chromosomes of the monitor lizard were mainly homologous to GGA28, with strong secondary sites at GGA31, GGA33 and Z that contained similar numbers of sex chromosome-borne genes (**Figure 4, Supplementary Fig. 2**).

### Candidate sex-determining genes of the four reptile species

Novel sex chromosomes may arise when an autosomal gene acquires a sex determining function. Sex chromosome turnovers have occurred many times during reptile evolution ^25, 26, 27^, possibly by a novel sex determining gene usurping the established gene ^28^ (e.g., *sdY* in rainbow trout ^29^, or a change to environmental sex determination and the subsequent evolution of novel genetic systems ^30^. It would not, therefore, be unexpected to find different candidate sex determining genes within the genomic regions that we have identified in the four reptiles (**Figures 1** and **4)**.

To test this, we compiled a list of genes reported to be involved in the sex-determining pathways of other vertebrates (**Supplementary Table S4, Supplementary Figs. S3-4**) and looked for their orthologs among genes that were either identified as sex-linked or fell within the identified sex-linked region in each studied species (**Figure 5A-B, Supplementary Table S5-7**).

**Figure 5.**
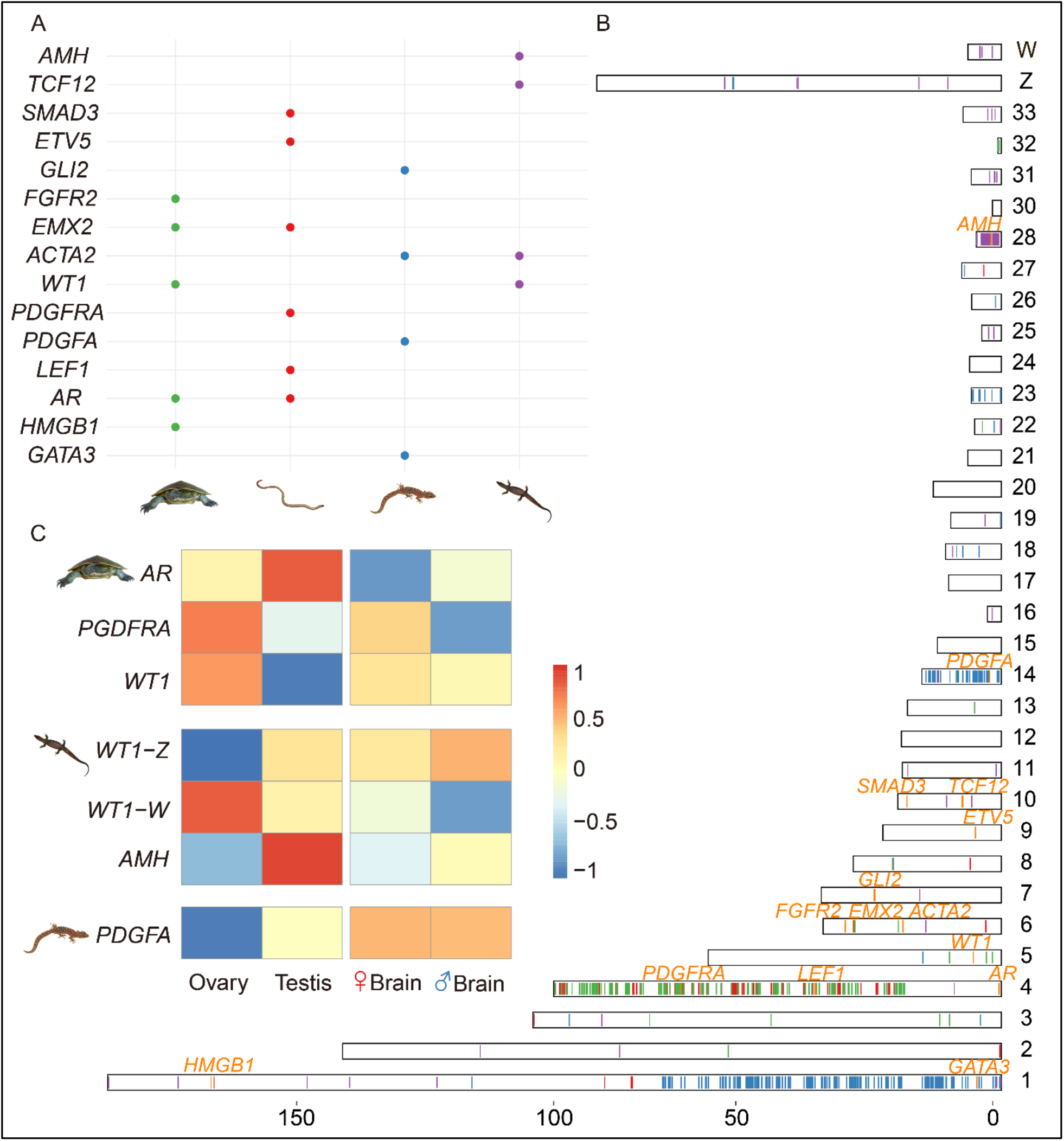
Candidate sex-determining genes of the four reptile species. **A**. The distribution of orthologs of vertebrate sex-determining genes that were also identified as on the sex chromosomes in this study. The coloured dots correspond to such genes within each species, which were identified by blast search against the chicken genome. For figures B, C and D, the river turtle is shown by green dots or bars, the monitor lizard by purple, the worm lizard by red and the marbled gecko by blue. B. Shows the ortholog positions of the sex chromosome-borne genes of these four reptile species on chicken chromosomes, with different colours of lines for different species’ orthologs. **C**. Gene expression patterns in the gonad and somatic tissues of candidate sex-determining genes of the three reptile species. We did not show it for the worm lizard due to unavailability of gonad tissues.

Included in the region on GGA4q that overlaps the homologous regions of the river turtle and the worm lizard X chromosomes, was one candidate male-determining gene *pdgfra* (platelet-derived growth factor receptor alpha). Another candidate sex determining gene *AR* (Androgen receptor) located on GGA4p was also annotated as X-borne in these two species. This suggests an independent acquisition of this gene because GGA4p is a microchromosome in all other birds which was fused recently. The *pdgfa* (platelet-derived growth factor alpha polypeptide) gene and its receptor *pdgfra* (platelet-derived growth factor receptor alpha) have been shown to be critical for testis development, particularly Leydig (male steroidogenic) cell development in mammals and turtles ^31, 32^, whereas *AR* is more likely to be involved in the downstream sexual differentiation process after the gonad sex is determined ^33, 34^. We confirmed *pdgfra* to be X-borne in the worm lizard using our transcriptome assembly and sex-linked probes from microdissected sex chromosomes. In the river turtle, we could not annotate *pdgfra* as a sex chromosome-borne gene because of a lack of mapped sex-chromosome borne probes, but it was embedded among other sex chromosome-borne genes in, so that is likely to be sex chromosome-borne also in the river turtle (**Figure 5B**).

For the marbled gecko, a promising candidate sex-determining gene is the Z-borne *pdgfa* (with a chicken orthologue on GGA14). For the monitor lizard, the most promising candidate sex-determining gene was *amh* (anti Mullerian hormone), which (or the duplicated copy of which) is located on GGA28 and plays a conserved role in testis development in multiple teleost species ^28^, birds ^35^, turtles ^36^ and even the platypus ^37^. Intriguingly, the ortholog of *wt1* (Wilms Tumour 1), an important regulator of *amh* and master male-determining gene *Sry* in human ^38^, was determined to be X and Y-borne in the river turtle, and Z and W-borne in the monitor lizard, and was located on a secondary chicken site of GGA5 (**Figure 5B, Supplementary Fig. 2**).

The expression patterns of these candidate sex-determining genes within the three reptiles for which we collected the gonad transcriptomes in this study further supported a function in the sex-determination pathway of each species (**Figure 5C, Supplementary Fig. S5**). The Z-borne *pdgfa* was specifically expressed in the testis of the marbled gecko; whereas its downstream receptor *pdgfra*, which is X-borne in the river turtle, was strongly expressed in the ovary. The X-borne *wt1* of the river turtle, as well as the W-linked *wt1* of the monitor lizard, were both expressed specifically in the ovary. The Z-borne *wt1* and *amh* of the monitor lizard were both specifically expressed in the testis. In summary, turnover of sex determining genes between the studied reptile species probably accounts for their sex chromosome turnovers.

### Evolution of dosage compensation and sex-linked gene expression in the four reptile species

Having identified the sex chromosome-borne genes of the four distantly related reptiles, we set out to examine their diversity of dosage compensation based on comparison of gene expression levels between sexes, and between the autosomes and the sex chromosomes. Since sex chromosomes may undergo meiotic sex inactivation in germ cells, and gonads are probably not appropriate for direct comparison between sexes ^39^, we focused on comparing the expression levels between sexes in their somatic (brain, tail or blood) tissues.

Among the four species, the worm lizard (with an XY sex system) and the monitor lizard (with a ZW sex system) have highly or moderately differentiated sex chromosomes. These two species exhibited a significantly (*P*<0.05, Wilcoxon test) different female vs. male expression ratio between autosomes and sex chromosomes (**Figure 6A, Supplementary Fig. S6**). The X-borne genes were more female-biased in the worm lizard, and Z-linked genes were more male-biased in the monitor lizard, indicating incomplete dosage compensation in the two species. In contrast, genes on the undifferentiated sex chromosomes, as well as autosomes, of marbled gecko and river turtle showed no significant difference of their expression ratios between sexes. This was because their Y- or W-linked genes have not degraded yet, so most genes on their X or Z chromosomes still have active partners on the Y (W) and thus there is no dosage difference between the sexes that selects for dosage compensation.

**Figure 6.**
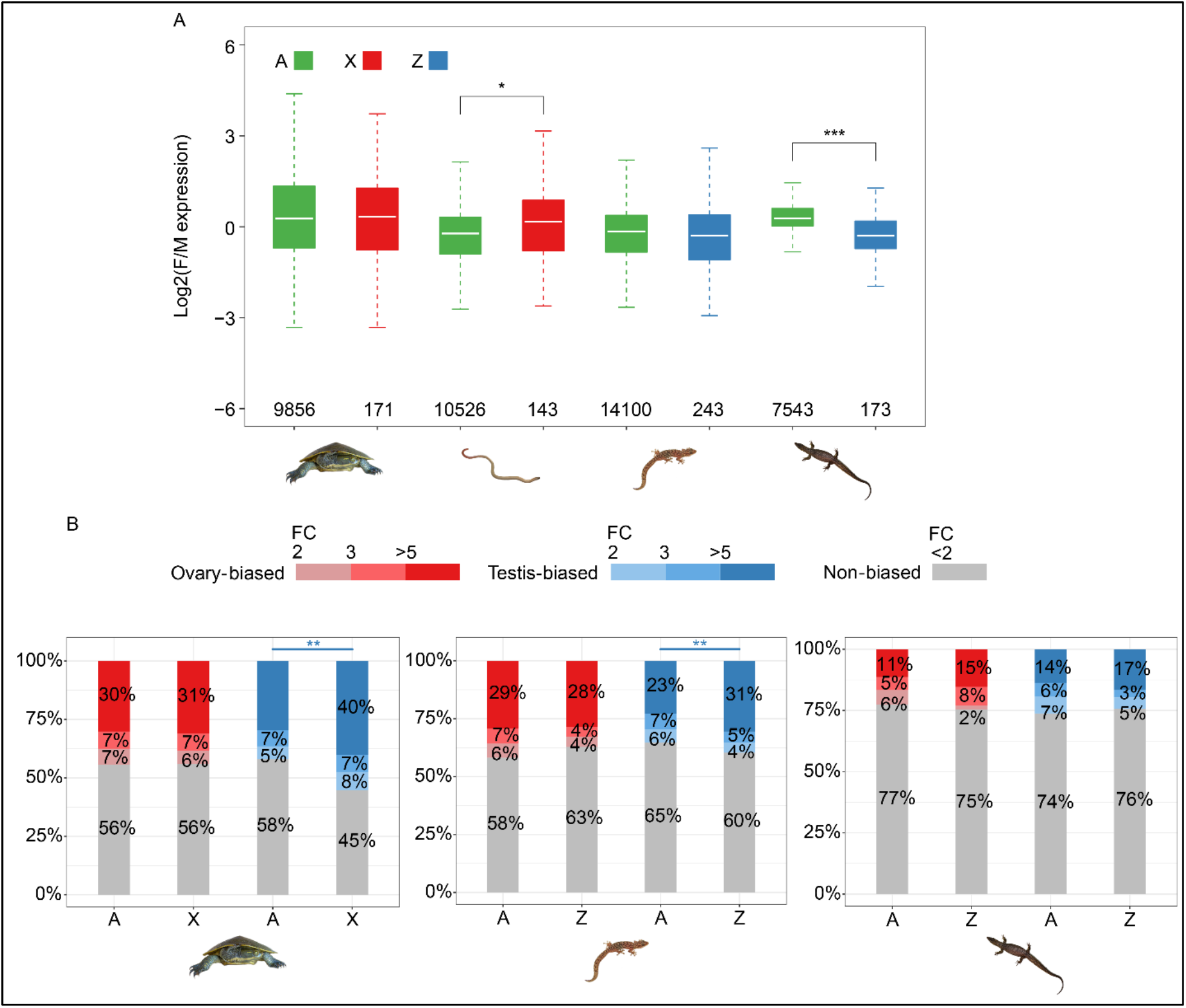
Dosage compensation and sex-linked gene expression in the four reptile species. **A**. Comparisons of the expression levels between sexes in the somatic (brain or tail) tissues of the four species. Genes from autosomes and the sex chromosomes are labelled in different colours, (autosomes: green, chrX: red and chrZ:blue). Only genes that have orthologs in chicken are considered and the respective gene numbers are shown on the X-axis. **B**. The comparisons of gonad specific gene expression levels between the sex chromosome against autosomes in three reptile species. Stacked bars shows the proportions of biased genes with more red or blue, the higher gonad-biased. Only genes with the most gonad-biased (the bluest/reddest part) were used in the significance testing when evaluating the level of masculinization.

For the three species with gonad transcriptomes (river turtle, marbled gecko and monitor lizard), we compared gene expression levels between sex chromosome vs. autosomes in the gonad, with the expectation that gonad-specific genes may have been preferentially selected to be located or not located on the sex chromosomes due to sex chromosomes’ sex-biased selective regimes. Previous studies in *Drosophila* ^40^ and other dipteran species ^41^ found underrepresentation of male-biased or testis-biased genes, and overrepresentation of female-biased or ovary-biased genes on the X chromosome, supporting such sex-biased selective regime. We found that testis-biased genes were overrepresented (*P*<0.001, chi-square test), while ovary-biased genes were underrepresented (*P*<0.005, chi-square test) on the Z chromosome of the marbled gecko (**Figure 6B**). However, a similar masculinization and defeminization pattern was not found on the undifferentiated Z chromosome of the monitor lizard, probably because few Z-borne genes were hemizygous (**Figure 4**).

The river turtle with undifferentiated XY sex chromosomes, unexpectedly showed a significant enrichment of testis-biased genes on the X chromosome relative to autosomal genes. This was probably because of cross-mapping of the reads of Y-borne genes that were not highly differentiated from those of X-borne genes. When we examined only the hemizygous X-linked genes (those without a Y-linked homolog) there was no such enrichment pattern. This suggests some Y-borne genes of the river turtle have undergone a masculinization process even though they were still retained by the Y chromosome.

## Discussion

Given the large genomes of many reptile species (up to 5.3Gb), fully sequencing sex chromosomes remains costly, despite the development of long-read sequencing and Hi-C technologies. So far, in depth studies of the gene content and dosage compensation of sex chromosomes have been carried out in a handful of lizards and snake species ^8, 42, 43, 44, 45, 46^ although ZW chromosome have been sequenced and a candidate sex determining gene identified in the central bearded dragon ^47^.

Here, we developed a cost-effective method to expand our knowledge of sex-linked genes and sex chromosomes in a range of non-model reptiles and applied it to four distantly related reptile species. We used it to map sex chromosome-borne genes from male and female transcriptomes that were identified by screening with DNA probes from microdissected sex chromosomes. We also applied the novel stLFR linked-read sequencing technology ^48^ and assembled the draft genome of monitor lizard, *V. acanthurus*, including the sex chromosome sequences. The newly identified gene content of the sex chromosomes of these four distantly related reptile species provided new insights into reptile sex chromosome evolution and dosage compensation.

Mapping the chicken orthologues of sex chromosome-borne genes of the monitor lizard (*V. acanthurus*), worm lizard (*A. parapulchella*) and marbled gecko (*C. marmoratus*) onto the chicken genome revealed examples of recruitment of different ancestral autosomes. We found that the sex chromosomes of the monitor lizard (*V. acanthurus*), worm lizard (*A. parapulchella*) and marbled gecko (*C. marmoratus*) have homologues on different chicken autosomes. This implies that they evolved from different autosomes in a common reptilian ancestor.

However, our finding that sex chromosomes of the distantly related pink-tailed worm lizard and river turtle (*E. macquarii*) both have homology to GGA4q provides a striking example of convergent recruitment of ancestral autosome regions. The long arm of the chicken chromosome 4 (GGA4q) has also been previously reported to be recruited as sex chromosomes of pygopodid gecko ^45^. This homology may signify that the same gene (likely to be *pdgfra*) has independently acquired a role in sex determination in all these species. Convergent recruitment of ancestral chromosome is a region orthologous to GGA23, which we identified to be part of the sex chromosomes of marbled gecko, and the central bearded dragon (*Pogona vitticeps*) ^46^.

Several general patterns emerged from these comparative analyses of the location of the chicken orthologues of genes on reptile sex chromosomes. Firstly, sex chromosomes seemed to have frequently originated by fusion of ancestral micro- and macro-chromosomes, or between micro-chromosomes ^49^. In addition to homology to the chicken microchromosome GGA28 that was reported, and also confirmed in this work as the ancestral sex chromosome of Anguimorpha species including spiny tailed monitor lizard ^50, 51^, we found that other chicken microchromosomes GGA31, 33 contained fragments homologous to genes on the sex chromosomes of spiny tailed monitor lizard. Microchromosomes also seemed to have contributed to the sex chromosomes of three other reptiles (**Figures 3 and 4**), as well as in the previously reported green anole lizard ^52^, bearded dragon lizard ^47^, soft-shell turtles ^53, 54^. The short arm of chicken chromosome 4 (which is homologous to the conserved region of the X chromosome of therian mammals, is a microchromosome in all species other than the Galliformes. These observations of homologies with chicken microchromosomes are not surprising given that half the chicken genes lie on microchromosomes.

A microchromosome origin might have contributed to the second feature of reptile sex chromosomes, most of which are less differentiated than those of birds and mammals. Homomorphic or partially differentiated sex chromosomes were found in three out of four reptiles we examined and are also described also in the giant musk turtle ^55^, eyelid geckos ^55, 56^ and some other gecko species ^57^, and skinks ^58^. The preponderance of poorly differentiated sex chromosomes in reptiles could be the result either of slow differentiation, or rapid turnover, or both. A potential cause for the generally slower rate of sex chromosome differentiation in reptiles could be the high recombination rate and gene density of the ancestral microchromosomes ^59^, which might prevent extensive recombination suppression and rapid differentiation between sex chromosomes in these reptiles.

Alternatively, rapid turnover of reptile sex chromosomes could explain the “ever young” partially differentiated sex chromosomes that are so common in reptiles. We have previously demonstrated ^5^ rapid transitions between sex determination systems in agamid lizards, and our present results expand the variety and independent origins of reptile sex chromosomes. In addition, the ability to switch into an environmental sex determination mode, and then to evolve novel genetic sex determination systems, may greatly facilitate turnovers. GSD and TSD have been reported within and between closely related reptile species, e.g., in agamid lizards ^60^, in viviparous skink ^61^, some turtles ^62^ and eye-lid geckos ^63^. In the Australian bearded dragon, the transition from GSD to TSD was observed both in the lab and in the field ^64^, despite its possession of a pair of highly differentiated sex microchromosomes ^65^.

Our identification of genes on reptile sex chromosomes enabled us to assess their transcription and assess dosage compensation. We found no evidence of global dosage compensation, even in the worm lizard *A. parapulchella* with highly differentiated X and Y chromosomes. This is similar to the absence of global dosage compensation in birds^66^ and other reptiles ^45^, but contrasts with the recently reported case of green anole lizard ^67, 68^, in which the single copy of the X chromosome is upregulated in XY males through an epigenetic mechanism similar to that in Drosophila. The absence of global dosage compensation in *A. parapulchella* could reflect dosage mitigation or tolerance at post-transcriptional levels, or it may be a consequence of its dosage-dependent sex-determination mechanism, similar to that in chicken, in contrast to a male-dominant XY system of the green anole.

In this work we combined cytogenetics and high-throughput sequencing to characterize the sex chromosomes of four reptile species. This greatly widened our knowledge of sex chromosome birth, death and dosage compensation in a vertebrate class that shows particular variety in modes and turnover of sex determining systems.

Thus, we used DNA from microdissected sex chromosomes to identify transcripts of genes located on the XY or ZW chromosome pairs in each species, and located their chicken orthologues on different chicken chromosomes. This revealed the diverse origins of sex chromosomes, but detected convergent evolution between distantly related reptiles (turtle and worm lizard). Our novel pipeline efficiently identified candidate sex determining genes, which differed from those of birds and mammals. We found that none of the four species showed transcription profiles expected of global chromosomal dosage compensation.

In summary, our molecular and cytogenetic characterisation of sex chromosomes in diverse taxa greatly expands our knowledge of reptile sex determination. By identifying reptile candidate sex genes and providing the means with which to identify more, we hope to realise the value of this particularly variable, but understudied, vertebrate class, the only one for which no master sex determining gene has yet been discovered.

The inexpensive and efficient method developed here can be applied to studying any species of eukaryote with cytologically distinct sex chromosomes, providing the basis with which to better understand the ecological and evolutionary drivers of sex chromosomes and sex determination systems.

## Materials and Methods

### Chromosome preparations, sex chromosome microdissection, probe preparations and FISH analysis

Animal collection, microdissection, preparation of sex chromosome specific probes and validation of probes were described in our previous studies ^18, 19, 20^. Briefly, we labelled sex chromosome probes by nick translation incorporating SpectrumGreen-dUTP (Abbott, North Chicago, Illinois, USA) or SpectrumOrange-dUTP (Abbott) and precipitated with 20 μg glycogen. After decantation, labeled probe pellets were resuspended in a 15 μl hybridization buffer. The resuspended probe mixture was hybridized with a drop of metaphase chromosome suspension fixed on a glass slide, covered with coverslips, and sealed with rubber cement. The slide was then denatured on a hot plate at 68.5°C for 5 min and was hybridized overnight in a humid chamber at 37°C for two days. The slides were then washed first with 0.4×SSC, 0.3% IGEPAL (Sigma-Aldrich) at 55°C for 2 min followed by 2×SSC, 0.1% IGEPAL for 1 min at room temperature. The slides were dehydrated by ethanol series and air-dried and then mounted with anti-fade medium Vectashield (Vector Laboratories, Burlingame, California, USA) containing 20 μg/ml DAPI (4′,6-diamidino-2-phenylindole.).

### Transcriptome assembly and annotation

RNA-Seq data from gonads and brain tissues for males and females of monitor lizard (*V. acanthurus*), river turtle (*E. macquarii*) and marbled gecko (*C. marmoratus*) and tail tissue from a male and female worm lizard (*A. parapulchella*) were used to perform *de novo* assembly of each species with Trinity v2.4.0 pipeline ^69^. Then we used transcoder ^69^ to do ORF prediction and cd-hit (v4.7) ^70^ to remove the redundant sequences with the parameters -c 1.00 -b 5 -T 8. For evaluating the quality of the assembly, we examine the number of transcripts that appear to be full-length or nearly full-length by BLAST+ (v2.6.0) ^71^ with the e-value 1e-3. For worm lizard and marbled gecko, the reference species is *G. japonicus* while for river turtle, the reference species is *P. sinensis*, and transcripts with a minimum 30% coverage of reference were selected. We used the Trinotate ^69^ pipeline to annotate the transcriptome. First, we aligned the transcripts to the reference library consisting of human and chicken using blastx and the protein file using blastp with the e-value 1e-3. Also, we used HMMER to do another annotation which aligned the transcripts to the Pfam protein library according to the hidden Markov algorithm with the default parameters. Later, the transcripts and the protein, along with the alignments from blast and HMMER were fed to Trinotate to annotate the transcriptome. The transcriptomes were evaluated by assessing the number of fully reconstructed coding transcripts with their reference species, which are *G. japonicus* for worm lizard and marbled gecko, and *P.sinensis* for river turtle.

### Genome assembly and annotation

We used SOAPdenovo2 (v2.0.4) pipeline ^72^ to assemble the Illumina DNA reads from microdissected sex chromosomes. In brief, we first tried several times with default parameters, to find the best K-mer with the longest N50. Then, we adjusted the average insertion size according to the best result and re-run the scaffold step. Afterwards, we used kgf(v1.16) with the parameters -m 5 -t 6 and Gapcloser(v1.12) to fill the gaps ^72^, which finally built a de novo draft assembly for sex chromosomes of our species.

We constructed the genome assembly of monitor lizard (*V. acanthurus*) with the Supernova v2.1.1 pipeline ^73^ with the default parameters, which is a package for de novo assembly based on 10X sequencing. Briefly, the approach is to first build an assembly using read kmers (default is 48), then resolve this assembly using read pairs (to K = 200), then use barcodes to effectively resolve this assembly to K ≈ 100,000. The final step pulls apart homologous chromosomes into phase blocks, which create diploid assemblies of large genomes.

We annotated the genome of monitor lizard (*V*. acanthurus) with the Braker2 v2.1.5 pipeline ^74^ which combined evidence of protein homology, transcriptome and *de novo* prediction. First, we used RepeatMasker (v4.0.7) ^75^ with parameters: -xsmall -species squamata -pa 40 -e ncbi, and the Repbase(v21.01) to annotate the repeat sequences. Then we aligned all available RNA-seq reads to the reference genome by STAR(v2.5) ^76^ to construct transcriptome evidence. Later we fed the masked genome, the alignment of RNA-seq, and the reference protein sequences, which were human and chicken here, to Braker2 with parameter: --prg=exonerate, setting exonerate for protein homology prediction. Finally, the package outputs the GFF file containing the gene models, along with protein sequences and CDS sequences. Additionally, we also separately annotated the sex chromosome of monitor lizard. First, we aligned our sequences to the reference protein sequences using tblastn with parameters: -F F -p tblastn -e 1e-5. The results were then refined by GeneWise (v2.4.1) ^77^, and for each candidate gene, we kept the one with the best score. Within these genes, we filtered them if premature stop codons or frameshift mutations reported by GeneWise or single-exon genes with a length shorter than 100bp, or multi-exon genes with a length shorter than 150bp, or if the repeat content of the CDS sequence is larger than 20% exists.

### Sex-linked sequences identification

For worm lizard (*A. parapulchella*), *river turtle* (*E. macquarii*) and marbled gecko (*C. marmoratus*), using an XY system as an example, we first assembled all the RNA-seq reads into a pooled transcriptome, the female reads into a XX transcriptome, and the male reads into a XY transcriptome. Then male RNA-seq reads were mapped to the XX transcriptome with bowtie2 (v2.2.9) ^78^ with default parameters. Read depth was then calculated using SAMtools (v1.6) ^79^, and those reads unmapped were assembled into a transcriptome which was considered to be X reads excluded.

The pooled transcriptomes were directly mapped by Illumina DNA reads from the X and Y chromosomes, and those sequences not mapped by either X or Y reads were assigned as autosomal genes. XX transcriptome and XY transcriptome were both mapped by DNA reads from the X and Y chromosomes, the reads depth (coverage/mappable site) was calculated for each genomic regions mapped, and sequences with a depth higher than 3 and a minimum coverage of 10% with X reads, simultaneously with no alignments with Y reads were assigned as X-linked. For the transcriptome with X reads excluded, the same steps were repeated and sequences with a depth higher than 3 and a minimum coverage of 10% with Y reads and no alignments with X reads were assigned as Y-linked. Afterwards, for sequences with both reads depth (X reads and Y reads) higher than 3, along with a minimum 10% coverage with both X and Y reads, we assigned them as shared genes.

To identify the sex-linked sequences in monitor lizard (*V. acanthurus*), Illumina reads from both sexes were aligned to the scaffold sequences using bowtie2 with default parameters. Read depth of each sex was then calculated using SAMtools in 10kb non-overlapping windows and normalized against the median value of depths per single base pair throughout the entire genome for the comparison between sexes. Those sequences with depth ratio of male-vs-female (M/F) ranging from 1.75 to 3, along with a read coverage ratio of male-vs-female higher than 0.8 were assigned as Z-linked sequences. For the rest of the sequences, those with M/F ratio of depth and coverage both ranging from 0.0 to 0.25 are assigned as W-linked sequences, and the remaining are assigned as autosomes.

### Homology comparisons

To find the orthologs of our genes with chicken, we compared the sex-linked transcripts of worm lizard (*A. parapulchella*), river turtle (*E. macquarii*) and marbled gecko (*C. marmoratus*), and the sex-linked genes annotated of the monitor lizard (*V. acanthurus*) 10X assembly to the proteins of chicken (v6, Ensembl), respectively using blastx with the e-value 1e-5. The result was filtered with the aligned AAs > 30% coverage of the reference chicken protein, along with a minimum 50% identity, and returned the one-to-one best hits, with the duplications retained. Then we merged the alignment sites from the four species and calculated the total number of orthologs on the relative chicken chromosomes. With the same protocols, we found the orthologs of our sex-linked genes with chicken sex-determining genes, except for the threshold of identity which were adjusted to 40%.

### Gene expression analyses

To quantify gene expression, RNA-seq reads were mapped to the transcripts of worm lizard (*A. parapulchella*), *river turtle* (*E. macquarii*) and marbled gecko (*C. marmoratus*) and the CDS of monitor lizard (*V. acanthurus*) by bowtie2. The raw read counts were estimated by RSEM (v1.3.1) ^80^, with both TPM expressions calculated. Those genes which have orthologs with chicken were filtered for dosage compensation analysis. Correlation within sexes for each species was tested using a Wilcoxon rank-sum test, where a significant differentiation within samples was found, with a p-value smaller than 0.05.

Gonadal biased genes were identified by calculating the fold change of gonadal to somatic expressions of the four species. For both ovary and testis, bias genes were classified into 4 categories of TPM ratio, namely <2, 2 to 3, 3 to 5, >5. For those genes that have a ratio <2 are assigned as negative, for those >=2, are assigned as gonadal biased, and only those genes with a ratio higher than 5 were calculated in the correlation test for masculinization.

### INDEL calling and PCR validation of sex specific markers

Using the identified 10.81 Mb Z-borne scaffolds and 7.10 Mb W-borne scaffolds of monitor lizard (*V. acanthurus*), we first identified indels based on their alignment using LASTZ(v1.02)^81^ with default parameters. On two homologous scaffolds (ChrZ_scaf_189 and Chr_W_scaf_176), we found three W-specific insertions with the lengths as 206 bp, 331 bp and 209 bp (**Supplementary Table S8**). At the flanking regions of these insertions, we designed two PCR primers spanning a predicted length of 1312 bp (assay 1) and 2479 bp (assay 2) sequence fragments for validating the sex chromosome specificity in monitor lizard (*V. acanthurus*). Both primer sets were amplified under the following conditions: initial denaturing at 95°C for 5 minutes followed by 35 cycles of 95°C for 30 seconds, 57°C for 30 seconds and an extension step of 72°C for 1 minute, with final extension of 72°C for 10 minutes. These two markers were validated on 60 individuals; 22 females and 38 males from 5 different localities distributed across the species distribution.

## Data Availability

The genomic and transcriptomic data of worm lizard (*A. parapulchella*), river turtle (*E. macquarii*), marbled gecko (*C. marmoratus*) and monitor lizard (*V. acanthurus*) have been deposited in GenBank under the BioProject accession code PRJNA737594. The draft genome and annotation of monitor lizard (*V. acanthurus*) have been deposited in Genome Warehouse (GWH) under the BioProject accession code PRJCA005583.

## Acknowledgments

Q.Z. is supported by the National Natural Science Foundation of China (32061130208, 31722050, 31671319), the Natural Science Foundation of Zhejiang Province (LD19C190001) and the European Research Council Starting Grant (grant agreement 677696). T.E. was supported by an ARC FT (FT110100733). This project was also partially supported by ARC DP (DP110102262) led by T.E. and University of Canberra Strategic Research Fund awarded to T.E. K.M. was supported by an ARC DP (DP110102262) led by T.E. T.G. was supported by NSF (IOS1146820). FS and JD were supported by the University of Canberra postgraduate research scholarships. Authors would like to acknowledge feedback by Janine Deakin on a preliminary draft. Animal photo credit: worm lizard and marbled gecko-TG; monitor lizard-JD; river turtle-AG and chicken- Liesl Taylor.

## Author contributions

TE, KM, JG conceived the idea. KM, FS, JD conducted lab works. ZZ and QZ conducted bioinformatic analyses. QZ, ZZ and TE wrote the first draft. All co-authors contributed intellectually to writing and editing the draft multiple times.

## Ethics Statement

Animal care and experimental procedures were performed following the guidelines of the Australian Capital Territory Animal Welfare Act 1992 (Section 40) and conducted under approval of the Committee for Ethics in Animal Experimentation at the University of Canberra (Permit Number: CEAE 11/07 and CEAE 11/12).

## Supplementary Figures Zhu et al.

**Supplementary Figure 1.**
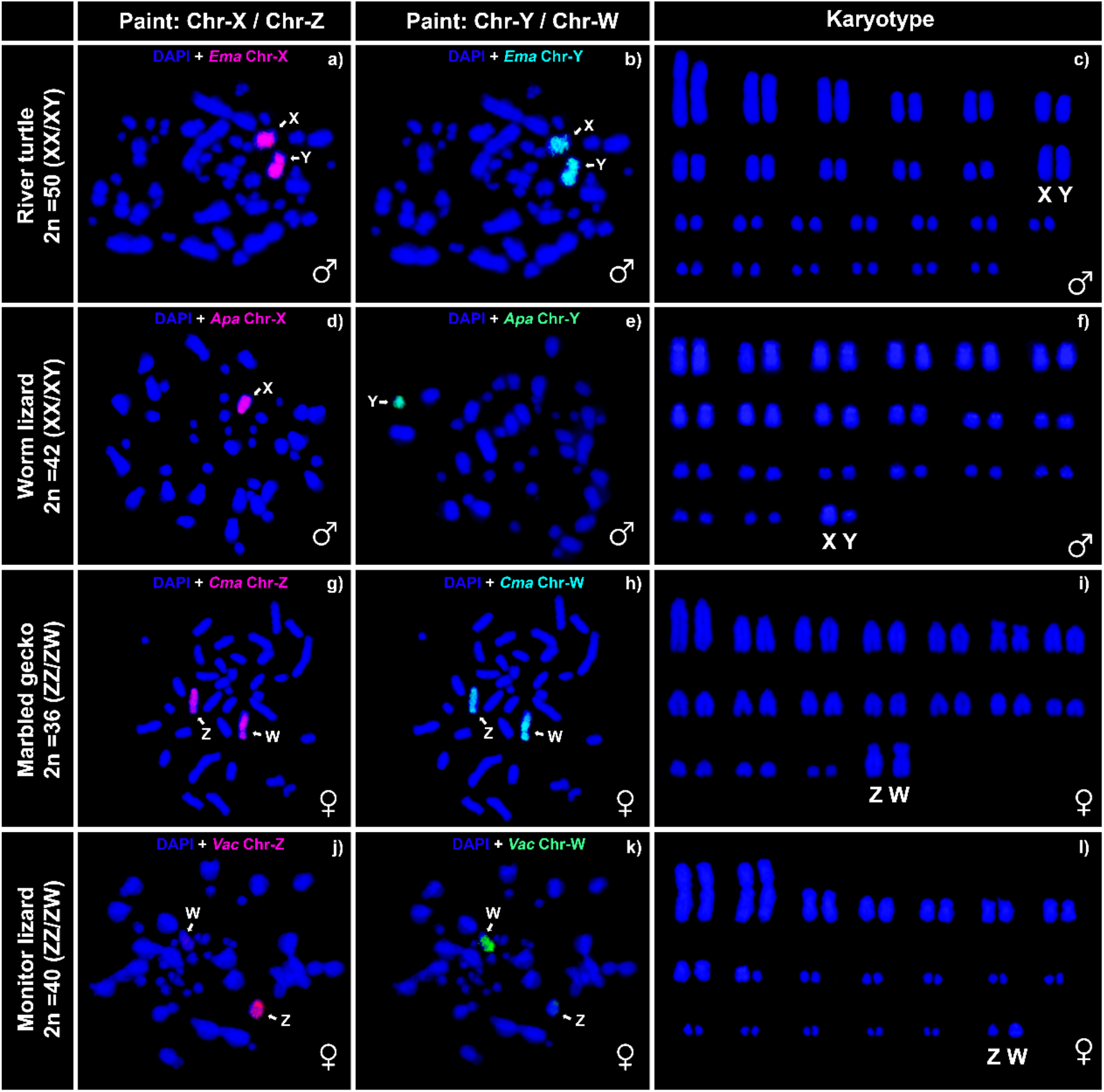
Florescence in situ hybridization (FISH) images of the four reptile species showing validation of microdissected chromosome probes.

**Supplementary Figure 2.**
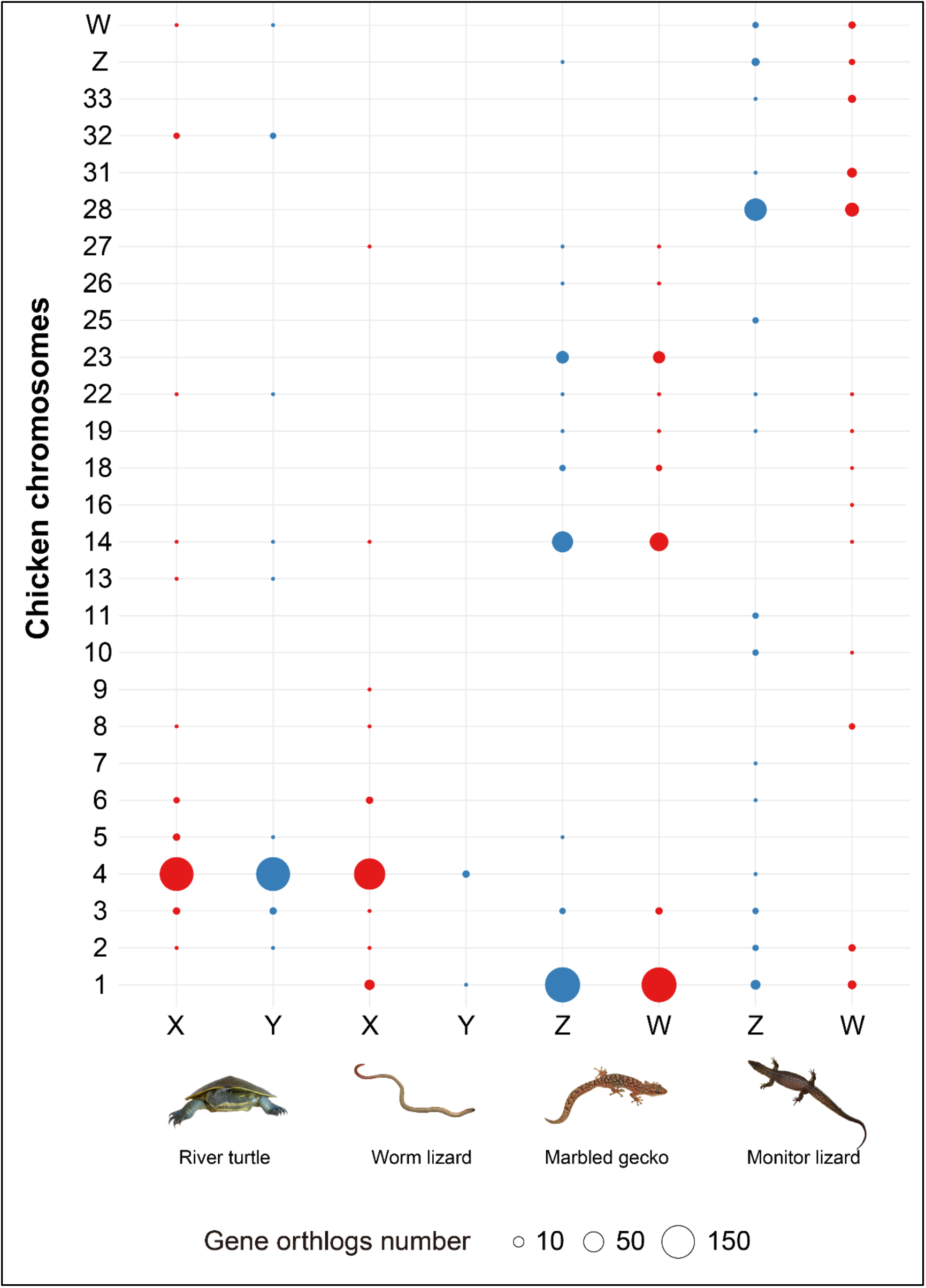
Bubbles showing orthologs of sex chromosomes of the four reptile species. The figure shows the origins of the sex chromosomes(X-axis) of four reptile species against the chicken genome(Y-axis). Dot sizes refers to the number of orthologs, and the color indicates the sex chromosomes of the four reptiles, which is red by X and W, and blue by Y and Z.

**Supplementary Figure 3.**
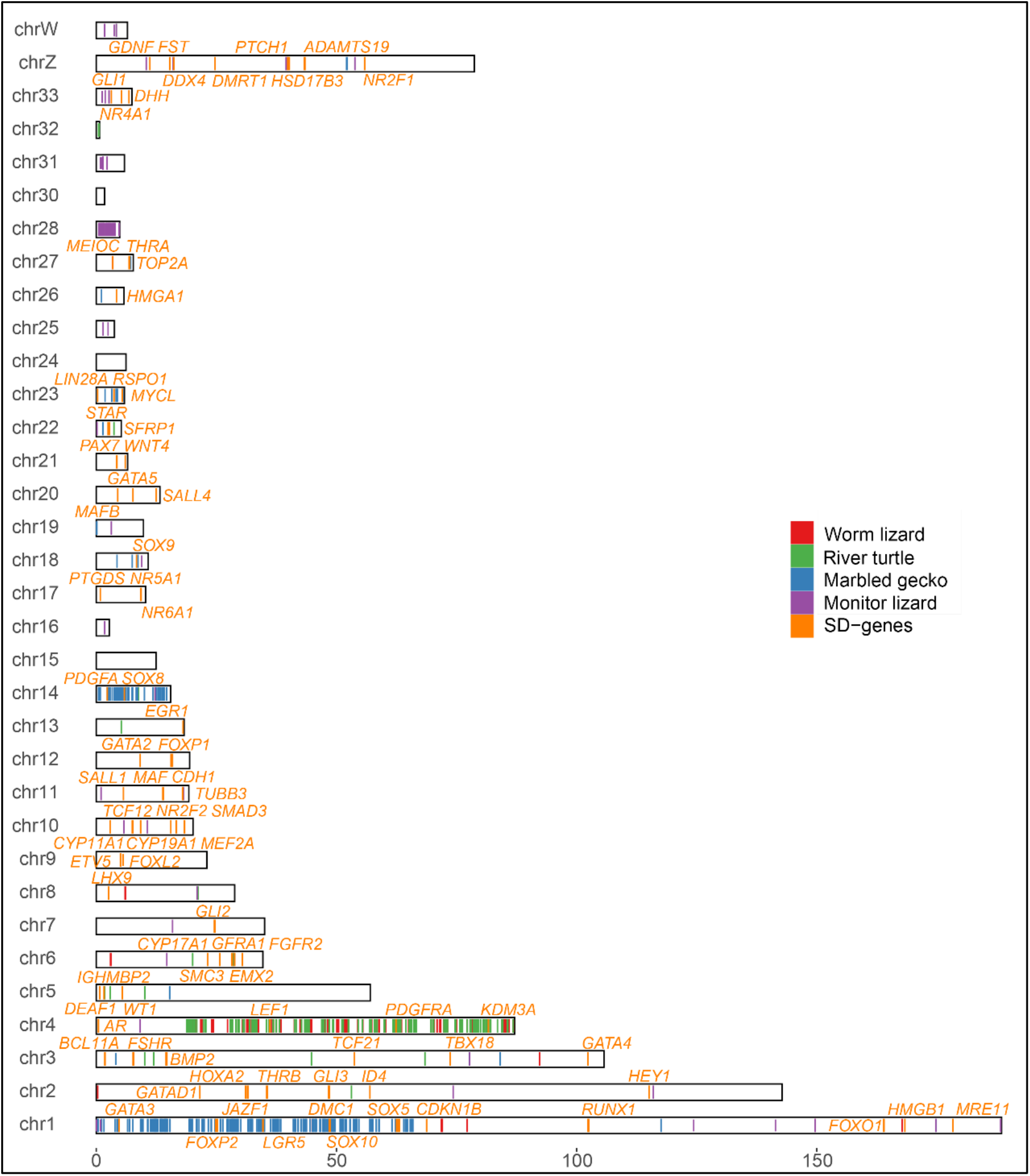
Location of orthologs of sex-determining genes on the homologous chicken chromosomes. We labelled the ortholog of known sex-determining genes in the chicken (Gga6a) genome. And different colors refer to genes of different species, with red to Worm lizard, green to River turtle, blue to Marbled gecko and purple to Monitor lizard. Lines in orange refers to the orthologs of sex-determining genes that have been studied. SD-genes: Candidate Sex determining genes

**Supplementary Figure 4.**
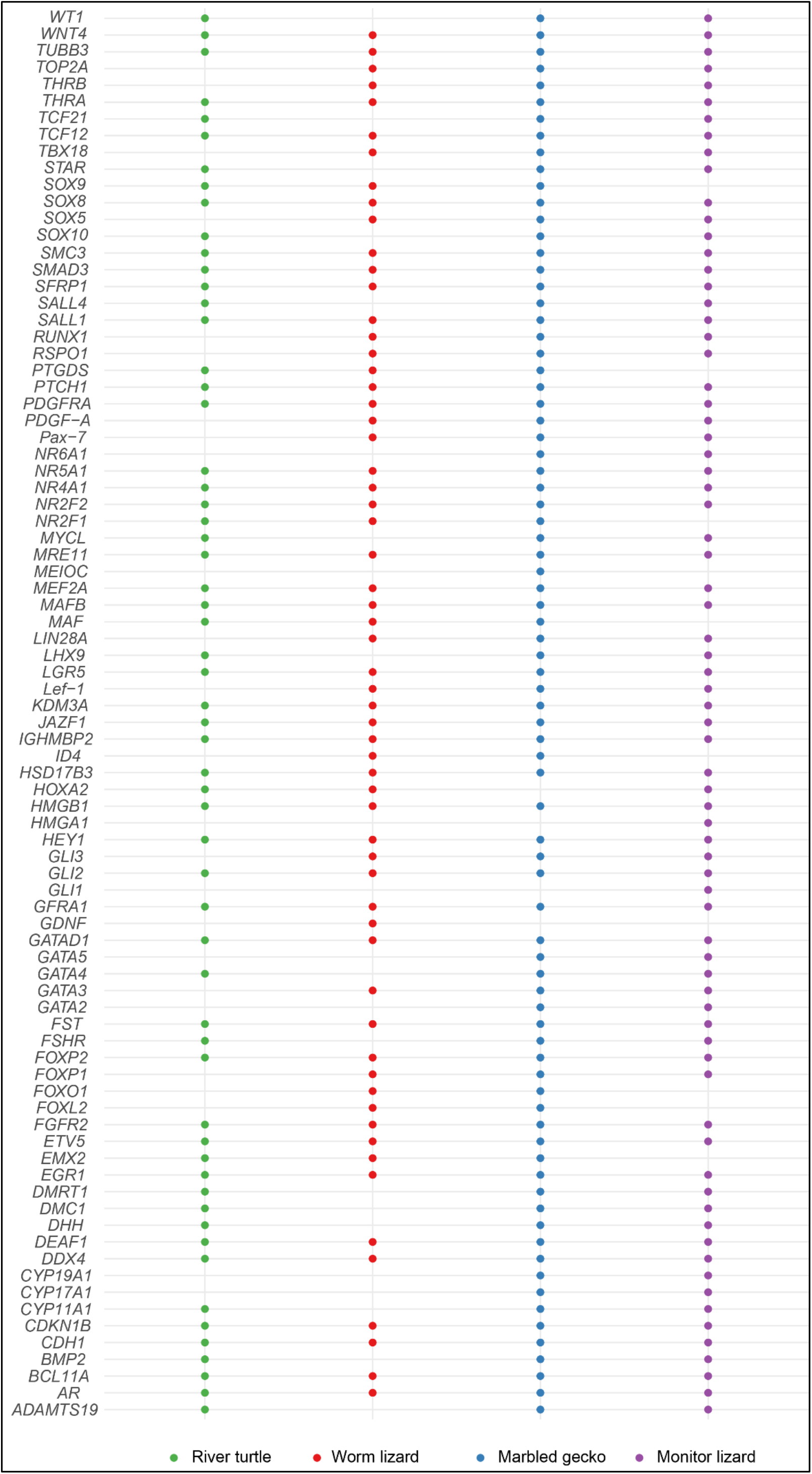
Orthologs of sex-determining genes ag. Gga6a in 4 species. The figure shows the presence (shown as a dot in the figure, otherwise no dot) of assembled orthologs of known sex-determining genes in the four species. And different colors refer to different species, which are red to Worm lizard, green to River turtle, blue to Marbled gecko and purple to Monitor lizard.

**Supplementary Figure 5.**
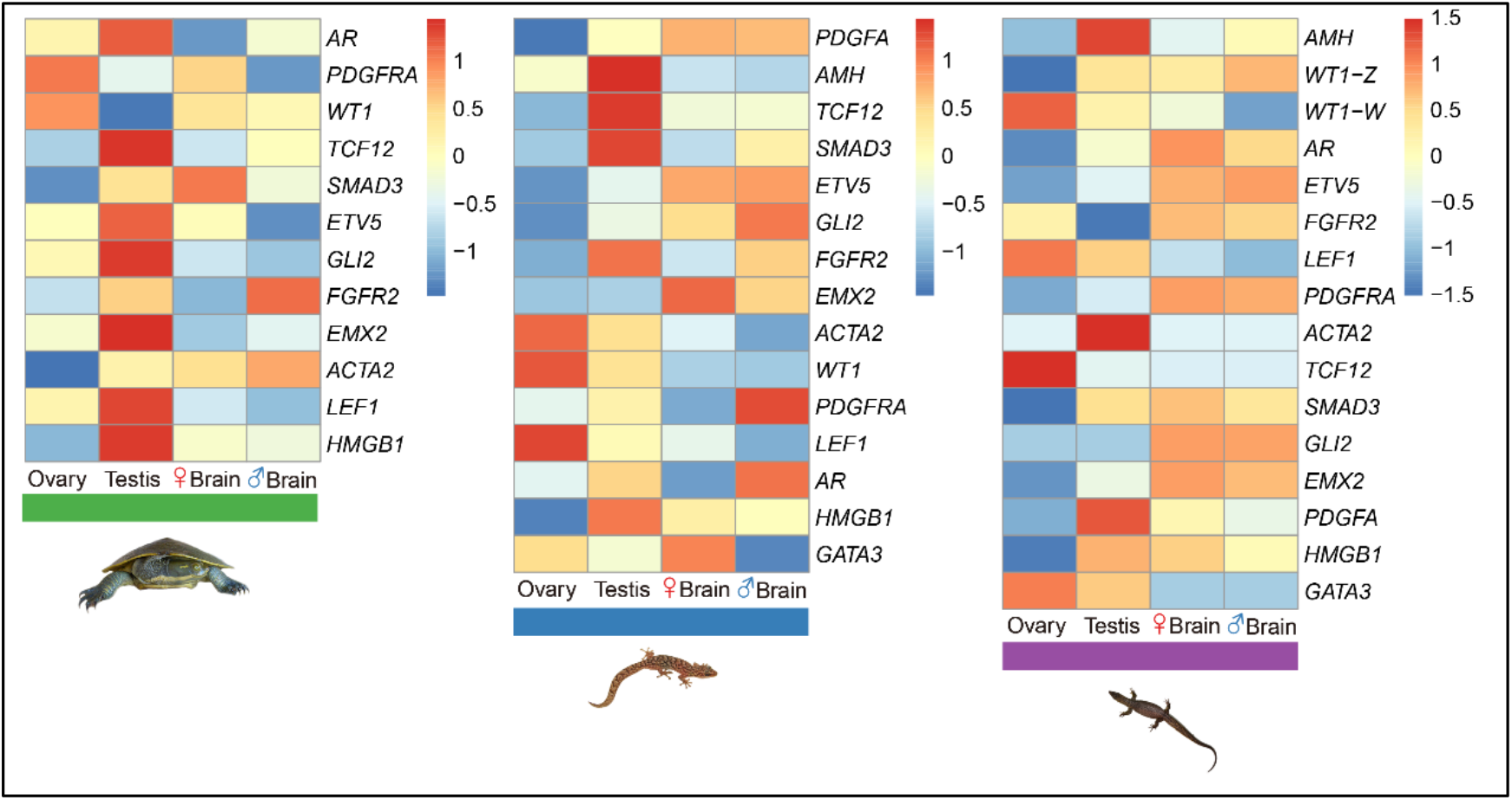
Comparisons of sex-determining genes’ expression patterns in different tissues. (Except for APA with only sexed somatic tissue transcriptomes.). The figure shows the log2 values of expressions (TPM) of assembled orthologs of known sex-determining genes in the three species. And different colors refer to different species, which are green to River turtle, blue to Marbled gecko and purple to Monitor lizard.

**Supplementary Figure 6.**
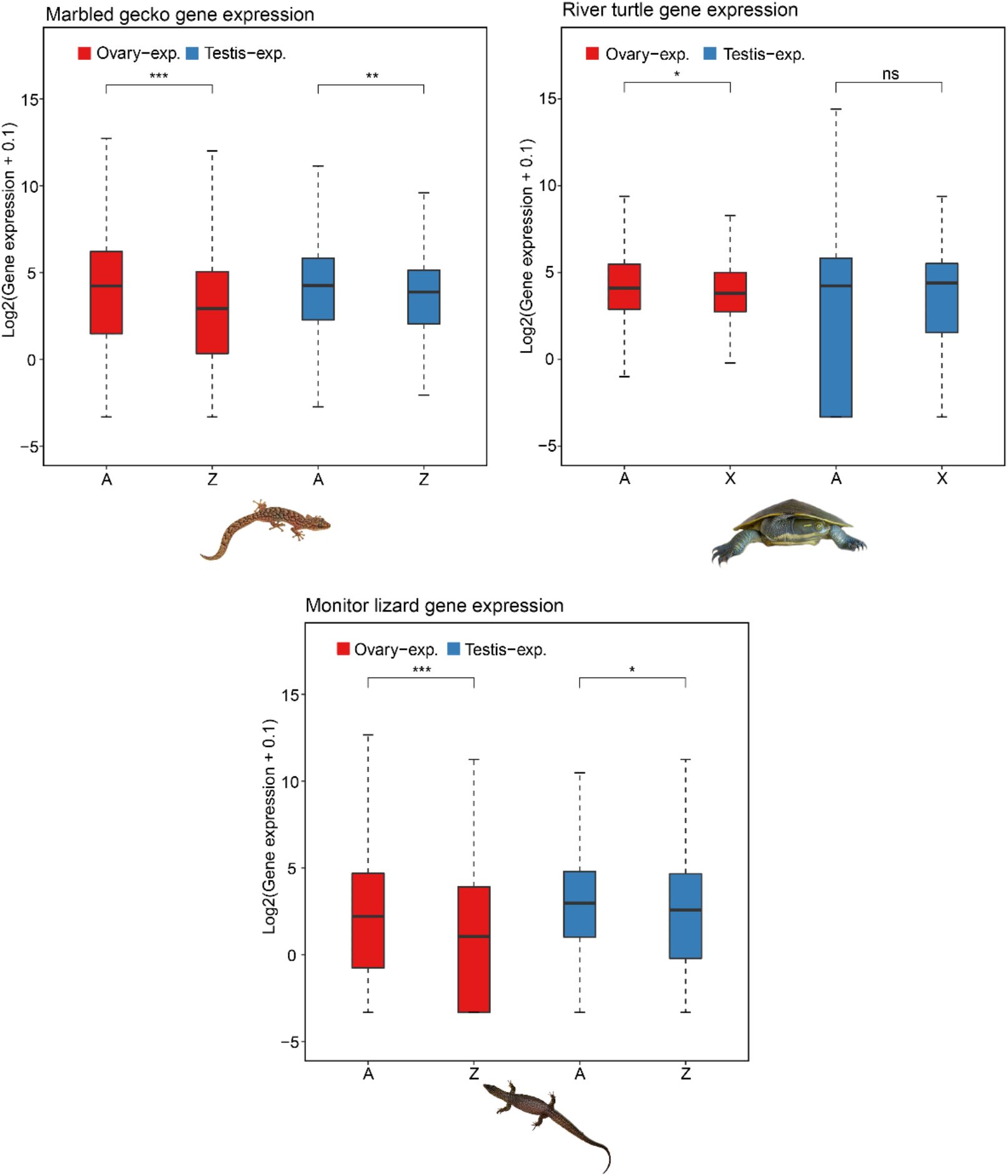
Gene expression of River turtle, Marbled gecko and Monitor lizard. Each box shows the log2 values of absolute gene expressions (TPM). A: autosomal genes; Z: Z-linked genes; X: X-linked genes. For XY species, to see masculinization, autosomal genes will generally have a higher testis expression. For ZW species, it is the opposite, and for ovary, it is all opposite.

